# Orangutan: an R package for analyzing and visualizing phenotypic data in the context of ecology and systematics

**DOI:** 10.64898/2025.12.18.695244

**Authors:** Javier Torres

## Abstract

**Aim:** Phenotypic characters have long been central to species diagnosis and delimitation and remain indispensable even in the age of genomics. However, phenotypic datasets are often complex— spanning dozens of traits of varying types and units, with correlated variables and unbalanced sampling—posing challenges for robust, reproducible analysis. Existing software solutions are fragmented, usually requiring labor-intensive workflows across multiple tools and manual steps, which undermines reproducibility and hinders comparisons across studies. To address these methodological and practical challenges, I introduce Orangutan, an R package designed to provide a flexible, easy-to-implement framework for comparing groups using mensural and meristic data.

**Innovation:** Orangutan provides a flexible and efficient framework for analyzing mensural and meristic data, supporting a full suite of statistical and visualization tools optimized for species delimitation and population comparisons. The package streamlines the identification of diagnostic, non-overlapping traits between species, while enabling rigorous assessment of both individual and multivariate trait differences. Core features include optional allometric correction to remove size effects, automated selection of appropriate univariate tests with post hoc comparisons, and integrated multivariate analyses. All outputs, including summary statistics and annotated publication-ready figures, are generated with minimal coding, ensuring accessibility and standardization.

**Main Conclusions:** Empirical validation with real-world datasets—including animal and plant species— demonstrates that Orangutan robustly identifies diagnostic traits, reveals both subtle and clear group differences, and achieves high classification accuracy with phenotypic data alone. By automating and unifying key analytical steps, Orangutan promotes reproducibility, transparency, and efficiency in phenotypic research. This package empowers researchers in taxonomy, ecology, and evolutionary biology to adopt quantitative best practices for species delimitation, facilitating comparative studies and advancing methodological standards in morphological data analysis. Orangutan is freely available with comprehensive documentation to support widespread adoption.

## INTRODUCTION

Phenotypic characters have long been the cornerstone of taxonomic practice, serving as the basis for species diagnoses and revealing observable boundaries between species (Balakrishnan, 2005). In fact, for practical and historical reasons most species have been described primarily on morphological grounds, even though genetic data are nowadays almost mandatory to delimit species. Phenotypic data remains indispensable for taxonomy, complementing molecular evidence and often revealing diagnostic differences that genetic data alone cannot capture (Cadena and Zapata, 2021).

Phenotypic datasets for ecology and systematics are inherently complex. They may include dozens of measurements per specimen and spanning diverse units (lengths, angles, indices, counts). In many cases trait values covary (e.g. allometric size effects), so one must normalize or apply allometric corrections to compare shape independently of body size. Data are often uneven—for example, few specimens per species but many characters or unbalanced group sizes. Small sample sizes combined with large numbers of variables invoke the “curse of dimensionality,” weakening statistical power and requiring feature selection or dimension-reduction (e.g. Principal Components Analysis—PCA). In practice, rigorous phenotypic analysis typically demands preprocessing (e.g. standardizing variables, applying size corrections), diagnostic trait discovery, and multivariate statistics to summarize variation.

Unfortunately, the software tools for phenotypic analyses are fragmented. Researchers often cobble together custom workflows using general statistics packages or multiple specialized tools. The result is laborious preprocessing (e.g. spreadsheet edits, separate script for each analysis), ad hoc statistical tests (Analysis of Variance, PCA, etc.), and piecemeal visualization—a process that weakens reproducibility and makes it difficult to compare results across studies. To my knowledge, there is no readily accessible, unified pipeline in R (or elsewhere) that integrates data cleaning, outlier handling, allometric adjustment, multivariate comparisons, and graphics specifically tailored to species delimitation with phenotypic data or more broadly, population comparisons.

Here I introduce *Orangutan*, an R package designed to fill this gap. *Orangutan* provides an end-to-end toolkit for phenotype-based species analysis: it uses a spreadsheet of mensural and meristic variables and identifies diagnostic traits—an important criterion for distinguishing species in the field. It also includes multivariate methods, such as PCA and Discriminant Analysis of Principal Components (DAPC), to explore the overall structure of phenotypic variation. For individual traits, *Orangutan* implements standard univariate statistical tests such as ANOVA or Kruskal–Wallis. All analyses are integrated with publication-ready visualizations, enabling users to efficiently generate informative and standard figures for their papers. By combining these steps into a reproducible workflow, *Orangutan* streamlines the process of comparing populations using phenotypic data, including delimiting and describing species.

In this manuscript I present *Orangutan*’s statistical and implementation framework in detail and demonstrate its utility with empirical case studies. I first review the methods and choices behind *Orangutan*’s analyses, then illustrate how the package can be applied to real taxonomic datasets. My goal is to empower researchers with a flexible, reproducible toolkit for phenotype-based species delimitation and to promote and democratize best practices in morphological data analysis.

## METHODS

*Orangutan* is an R package designed to facilitate the statistical analysis and visualization of phenotypic data in the context of species descriptions or, in general, population comparisons. The package streamlines phenotypic data handling, allowing users to load structured datasets, conduct statistical comparisons, and generate informative visualizations with minimal coding effort (e.g. full execution in a single function—*run_orangutan*). All visualizations are saved with publication quality, e.g., TIFF format, 7.5 in. width, 6 in. height, and 300 dpi.

*Orangutan* supports a multi-faceted approach to assessing phenotypic distinctiveness between groups. First, it enables the identification of non-overlapping traits—traits whose distributions do not overlap between groups—offering a clear and often desirable form of phenotypic diagnosability when delimiting species (Braby et al., 2024). Second, the package allows users to explore differentiation based on entire suites of traits by conducting multivariate analyses, including clustering and classification. These analyses assess whether groups form distinct clusters in multivariate space and whether individuals can be accurately assigned to their respective groups based on phenotypic characters alone. Third, *Orangutan* facilitates the detection of statistically significant differences in individual trait values, even when their distributions overlap. Such statistically significant differences provide evidence of consistent, average-level divergence that may indicate underlying genetic, ecological, or evolutionary separation.

Key functionalities include identifying and removing outliers (optionally), detecting non-overlapping traits, applying allometric corrections before multivariate analysis (optionally), and automatically selecting and conducting appropriate parametric or non-parametric tests for univariate analyses. By integrating these components, *Orangutan* enables researchers to robustly and reproducibly analyze phenotypic variation, assess different dimensions of phenotypic distinctness, and identify traits relevant to species delimitation with ease. The package is freely available at https://github.com/metalofis/Orangutan-R with detailed instructions for installation and execution.

### Dependencies and input data

The *Orangutan* package depends on a suite of R libraries for statistical analysis, data manipulation, and visualization. These include *adegenet* (Jombart, 2008), used for multivariate analysis, specifically implementing DAPC. *dunn*.*test* (Dinno, 2024) performs post-hoc pairwise comparisons following Kruskal-Wallis tests to identify significant differences among species. *dplyr* (Wickham et al., 2023) facilitates data manipulation tasks such as filtering, selecting, sorting, and summarizing datasets, supporting efficient data cleaning and transformation. *ggplot2* and *ggpubr* (Kassambara, 2023; Wickham, 2016) create layered visualizations including PCA plots, DAPC plots, and boxplots. *multcompView* converts post-hoc test results into letter-based groupings that can be overlaid on plots to highlight significant differences. *tidyr* (Wickham and Girlich, 2022) reshapes data for analysis and plotting, ensuring datasets are in the correct format. RColorBrewer (Neuwirth, 2002) supplies distinct color palettes, improving the interpretability of species-specific plots.

As data input, *Orangutan* expects phenotypic data in a structured format as a comma-separated values (CSV) file where the first column—species—contains species designations, and the remaining columns contain meristic and mensural variables, including one column for the main_length, which is optionally used for allometric correction (explained in “Allometric Transformation of Mensural Traits”). Upon loading, the data undergoes removal of unwanted columns, such as generic index columns, and sorting alphabetically by species names to ensure consistency across downstream analyses. To further enhance the user experience, *Orangutan* assigns colors to species dynamically using the RColorBrewer palettes, with the number of colors tailored to the number of species in the dataset. If the number of species exceeds the palette’s default limit (e.g., 12 colors for Paired), the color scheme is extended by cycling through available colors.

I used three empirical datasets to validate *Orangutan* by running them to test the different elements of the package. One dataset comprises four species of anole lizards and one putative hybrid category (Torres, 2024). Another dataset comprises nine species of anole lizards (Torres et al., 2025). A third dataset is the “Iris” dataset included in R. It comprises three plant species and the main difference from the anole datasets is that the “Iris” dataset does not require allometric correction. The anole datasets can be downloaded from https://github.com/metalofis/Orangutan-R/tree/main/example_datasets.

### Summary Statistics

*Orangutan* computes summary statistics for each numeric trait by species, enabling reproducible and straightforward analysis. It automatically identifies numeric variables, adapting to different datasets without requiring user input. For each trait, it calculates the mean ± standard deviation, along with minimum and maximum values, summarizing results as “Mean ± SD (Min - Max).” Outputs are exported as a CSV file for easy reporting.

### Identification of Non-Overlapping Variables

To evaluate species diagnosis, *Orangutan* identifies phenotypic traits with non-overlapping value ranges between species pairs—useful for diagnosis. It checks whether a trait’s range in one species does not intersect with that of another. When found, non-overlapping traits are visualized with boxplots using consistent species colors. To report values, users can simply check the summary statistics table for the variable and species pairs that do not overlap. If overlapping traits are not found, a message is provided. The overlapping traits are later tested in univariate analyses for statistical differences.

### Multivariate Analyses

#### Allometric Transformation of Mensural Traits

To ensure that trait comparisons are biologically meaningful, *Orangutan* includes an optional allometric correction step using the “main length” of the studied organism as a size proxy. The main length represents a standard body or structure size against which other measurements are scaled. For animals, this might be snout–vent length in amphibians and reptiles, or total body length in fish. For plants, it could be stem height, or inflorescence axis length, depending on the structure being analyzed. If allometric correction is not needed, the input dataset does not need a main_lengh column.

This step is essential for distinguishing shape-based differences from size-related effects, particularly when comparing mensural traits. The allometric adjustment process begins by identifying the column representing “main length” and selecting mensural variables. Both the main length and the rest of the mensural variables are then log-transformed to linearize the allometric relationships. A linear regression is performed for each mensural trait against log-transformed main_length, and the residuals from this regression represent size-independent trait variation (Nakagawa et al., 2017). The final dataset consists of these residuals, which are combined with species classification, while non-mensural traits remain unchanged. This allometric transformation ensures that subsequent statistical analyses focus on shape differences rather than size variation, improving the reliability of species comparisons.

#### Principal Component Analysis (PCA)

To assess whether candidate species form distinct clusters in multivariate space, a PCA is performed only with numeric values. The PCA is conducted using the *prcomp* R function, with the data centered and scaled for standardizing (Bro and Smilde, 2003). The first two principal components (PC1 and PC2) are extracted for further analysis. The contribution of each variable to PC1 is assessed by calculating the absolute values of the loading scores, and the top nine variables contributing most to PC1 and PC2 are identified. The explained variance for each principal component is computed by squaring the standard deviations of the principal components and dividing by the total variance, which is then expressed as a percentage on the plot. For visualization, polygons are created for any species declared by the user to delineate their phenotypic space.

#### Discriminant Analysis of Principal Components (DAPC)

DAPC is conducted also to examine group clustering patterns. Using the *dapc* function from the *adegenet* package, the analysis retains 10 principal components and 5 discriminant axes (LDs). The jackknife method is applied to assess classification robustness and estimate the error rate. The coordinates of each individual along the first two linear discriminants (LD1 and LD2) are extracted and stored in a dataframe. The proportion of variance explained by LD1 and LD2 is calculated by dividing the corresponding eigenvalues by the sum of all eigenvalues. Similar to PCA, polygons can be overlaid on the DAPC scatter plot for any species.

To assess classification performance, a confusion matrix is generated by comparing true species labels with predicted group assignments (Table 2). This matrix summarizes the results of jackknife-resampled cross-classification from the linear discriminant analysis. Rows show the *a priori* classifications, while columns show the model’s predicted classifications. Diagonal elements indicate correctly classified individuals, and off-diagonal elements indicate misclassifications. Elements above the diagonal represent false positives for the column class— cases where the model incorrectly assigned individuals from a given actual class to a different predicted class. Elements below the diagonal represent false negatives for the row class—cases where the model predicted individuals belonged to a class when they belonged to the row’s class.

### Univariate Analyses

#### ANOVA, Kruskal-Wallis and post hoc tests

To assess phenotypic trait differences across species/populations, one-way Analysis of Variance (ANOVA) is performed for each variable. The assumptions of normality and homogeneity of variances are tested using the Shapiro-Wilk test and Bartlett’s test, respectively. If both assumptions are met (p-values > 0.05), ANOVA is conducted, and the results are interpreted based on the F-statistic, degrees of freedom, and p-value. For variables with significant ANOVA results (p-value < 0.05), post-hoc pairwise comparisons are performed using Tukey’s Honest Significant Difference (HSD) test. The results are summarized and visualized with boxplots, which are annotated with Tukey HSD labels to highlight significant differences. For variables failing ANOVA assumptions, the non-parametric Kruskal-Wallis test is used to compare traits between species. If the Kruskal-Wallis test is significant (p-value < 0.05), pairwise post-hoc comparisons are performed using Dunn’s test with a Bonferroni correction. The results are summarized in a table and visualized with boxplots annotated with Dunn’s test labels.

#### Summary Tables

Univariate analyses results are saved in summary tables. The ANOVA summary table reports, for each variable, the F-statistic, p-value, and whether the assumptions of normality and homogeneity of variances were met. The Kruskal-Wallis summary table reports, for each variable, the p-value, Chi-squared statistic, and any significant pairwise differences detected by Dunn’s post hoc test. These tables enable a clear overview of which variables show significant differences among groups and by which statistical method the significance was determined.

## RESULTS

I successfully ran *Orangutan* on the three empirical datasets and below I show output examples from all analyses.

### Summary Statistics

The function *summary_stats* produces a CSV file with summary statistics for each species or population (Table 1), which is a common report in species descriptions to have a first, general idea of the variation among populations and whether there are differences amongst them. This function will dynamically identified numeric variables and ran the summary statistics on only those when the dataset also contained categorical variables.

**Table 1.** Summary statistics (mean ± standard deviation and range) of a partial dataset from Torres et al. 2025. A: *Anolis*.

| Species | Snout-vent length | Head length | Postmentals |
| --- | --- | --- | --- |
| *A. allisoni* | 82.36 ± 7.1 (73–94.7) | 26.48 ± 2.33 (24.7–31.8) | 5.89 ± 0.78 (4–7) |
| *A. brunneus* | 70.08 ± 1.42 (67–71.9) | 21.22 ± 1.1 (19.7–23) | 6.11 ± 0.33 (6–7) |
| *A. carolinensis* | 59.3 ± 3.23 (55–63.3) | 17.16 ± 1.31 (15.6–19.17) | 6.12 ± 0.83 (5–8) |
| *A. fairchildi* | 68.58 ± 1.85 (66–70.3) | 20.86 ± 1.78 (18.9–22.4) | 5.5 ± 1 (4–6) |
| *A. longiceps* | 72.87 ± 4.42 (64.96–78.84) | 22.96 ± 2.33 (20–27.21) | 4.14 ± 0.38 (4–5) |
| *A. maynardii* | 69.3 ± 0.71 (68.8–69.8) | 21.5 ± 0.14 (21.4–21.6) | 7 ± 1.41 (6–8) |
| *A. porcatus* | 66.3 ± 8.03 (59.8–77.9) | 21.45 ± 2.87 (18.79–25.4) | 6 ± 0 (6–6) |
| *A. smaragdinus* | 59.31 ± 3.66 (52.6–64.7) | 16.19 ± 1.65 (13.3–18.9) | 4.9 ± 1.2 (4–7) |
| *A. torresfundorai* | 70.14 ± 5.19 (62.8–78) | 22.42 ± 2.39 (18.7–26.08) | 5.89 ± 1.05 (4–8) |

**Table 2.** Cross-classification table resulting from the Torres et al. (2025) dataset: Jackknife-resampled cross-classification resulting from the linear discriminant analysis. Rows represent a priori classification, while columns represent the model’s classification. Values in the diagonal are correctly classified individuals, whereas values outside it are misclassifications, for an accuracy of 71.3% (57 of 80 individuals correctly classified). The elements above the diagonal are instances in which a sample from a row’s actual class was incorrectly classified as a class to the right (a different predicted class). These are the false positives for the column class. The elements below the diagonal represent instances in which a sample from a column’s predicted class was incorrectly classified as the class corresponding to the row. These are the false negatives for the row class. Species are *Anolis allisoni* (all), *A. brunneus* (bru), *A. carolinensis* (car), *A. fairchildi* (fai), *A. longiceps* (lon), *A. maynardii* (may), *A. porcatus* (por), *A. smaragdinus* (sma), and *A. torresfundorai* (tor).

|  | alli | bru | car | fai | lon | may | por | sma | tor |
| --- | --- | --- | --- | --- | --- | --- | --- | --- | --- |
| alli | 9 | 1 | 0 | 0 | 0 | 0 | 0 | 0 | 0 |
| bru | 0 | 8 | 0 | 1 | 0 | 0 | 1 | 0 | 0 |
| car | 0 | 0 | 9 | 0 | 0 | 0 | 0 | 0 | 0 |
| fai | 0 | 0 | 0 | 4 | 0 | 0 | 0 | 1 | 1 |
| lon | 0 | 0 | 0 | 0 | 10 | 0 | 0 | 0 | 0 |
| may | 0 | 0 | 1 | 0 | 0 | 2 | 0 | 0 | 0 |
| por | 0 | 0 | 0 | 0 | 0 | 2 | 6 | 2 | 2 |
| sma | 0 | 0 | 0 | 1 | 0 | 0 | 2 | 6 | 1 |
| tor | 0 | 1 | 0 | 0 | 0 | 0 | 3 | 0 | 6 |

### Identification of Non-Overlapping Variables

*Orangutan* identified variables that were diagnostic across variables and species pairs and produced one plot per variable per species pair (Fig. 1). The colors were automatically assigned to each species/population. The plots produced visually summarize the distribution of the dataset using five key statistics (Fig. 1). The box represents the interquartile range (IQR), spanning from the first quartile (25th percentile) to the third quartile (75th percentile), which contains the middle 50% of the data. A horizontal line inside the box marks the median, indicating the central tendency. Whiskers extend from the box to the smallest and largest values within 1.5 times the IQR from the lower and upper quartiles, respectively. Data points beyond this range are plotted individually as outliers, highlighting unusually high or low values in the dataset.

**Figure 1.**
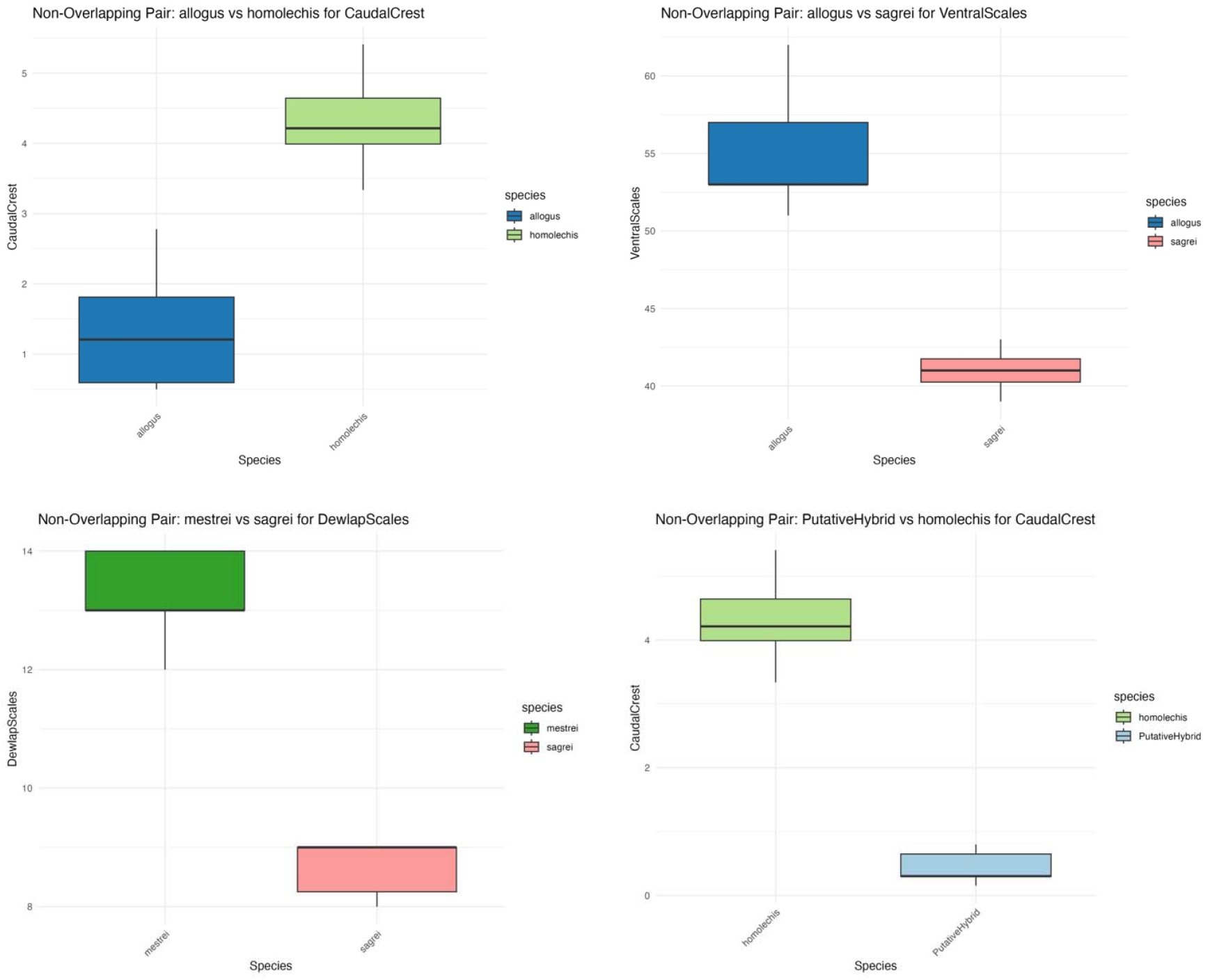
Sample of four of the non-overlapping variables identified by *Orangutan* from the anole dataset from Torres (2024).

### Multivariate Analyses

The multivariate analyses conducted PCA and DAPC and produced plots (Fig. 2). A confusion matrix from the DAPC classification is also produced, showcasing the accuracy of species assignments (Table 2).

**Figure 2.**
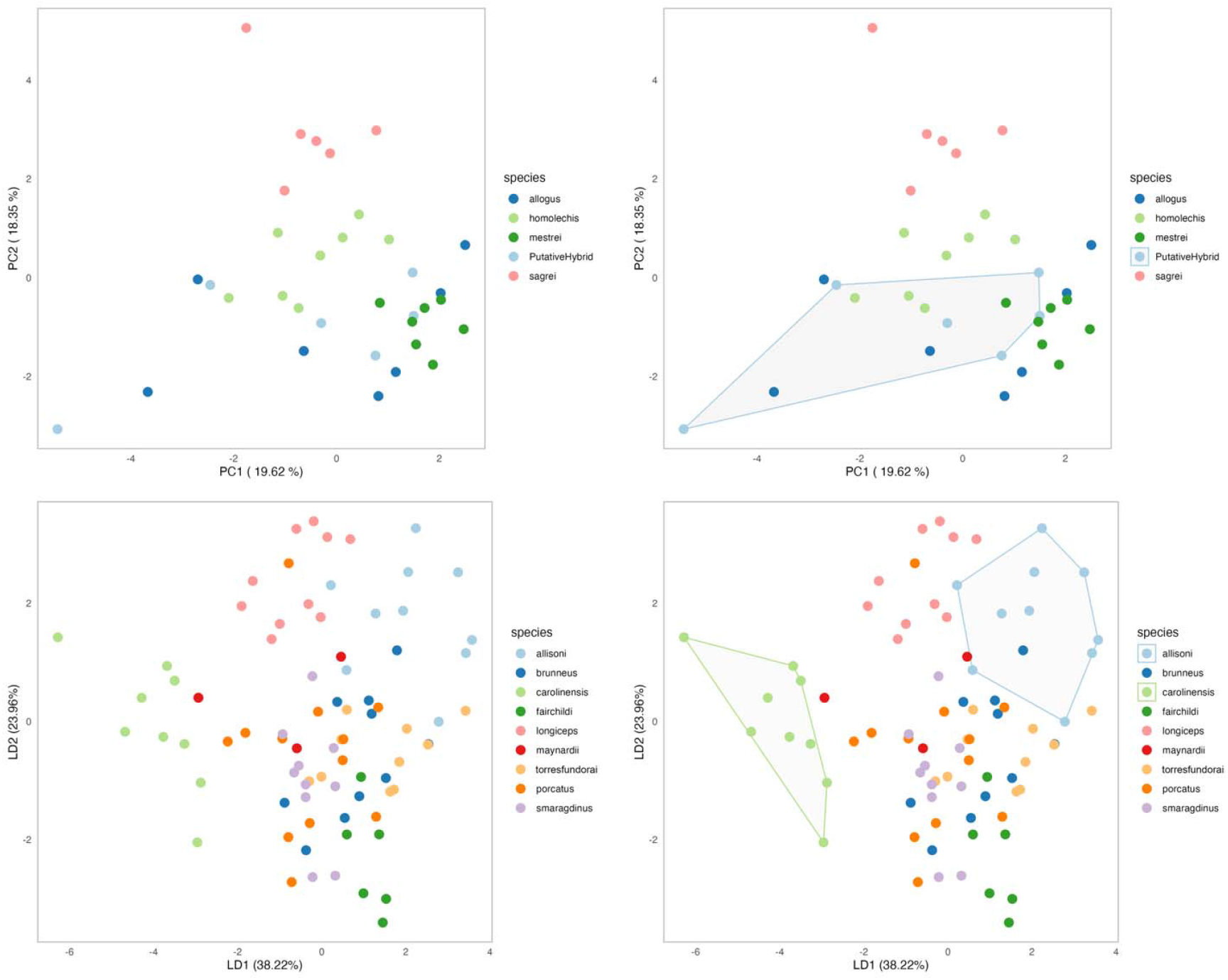
Multivariate analyses visualizations. PCA on the top using the Torres (2024) dataset. DAPC on the bottom using the Torres et al. (2025) dataset. The right panel depicts the option of declaring species to encircle, useful for emphasizing a given species.

### Univariate Analyses

Univariate analysis identified significant differences among species for several variables across the three datasets. From a total of 31 variables, 14 met the assumptions of normality (Shapiro-Wilk test) and homogeneity of variances (Bartlett’s test) and were analyzed using ANOVA. Among these, 13 variables showed significant differences (p < 0.05). Post-hoc Tukey’s Honest Significant Difference (HSD) tests revealed distinct groupings among species. The results, including F-statistics, degrees of freedom, and p-values, were summarized in summary_anova.csv. Corresponding boxplots annotated with Tukey’s HSD groupings were generated for visualization (e.g., Fig. 3).

**Figure 3.**
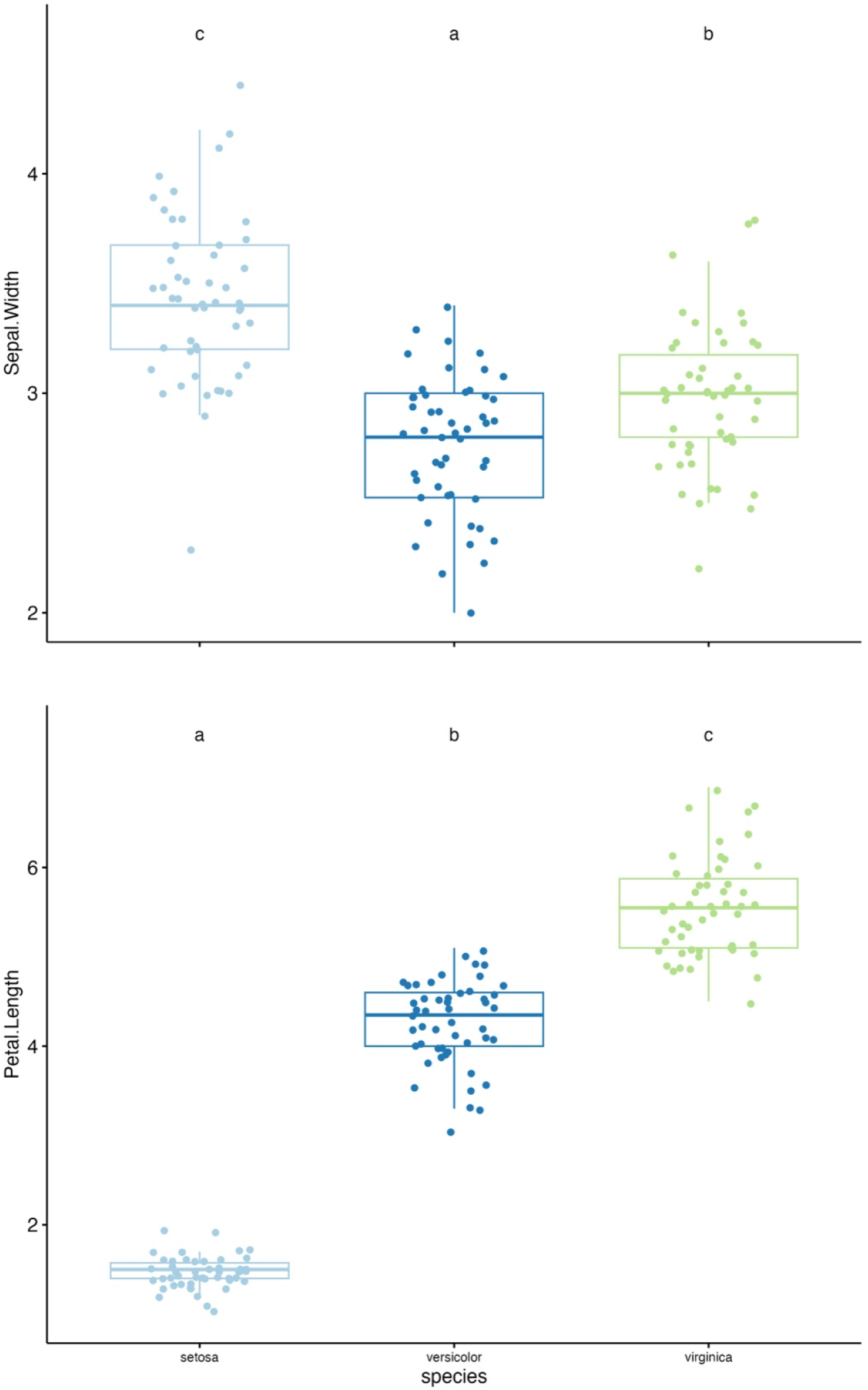
Univariate visualizations using the R “Iris” dataset. Horizontal line = median, box = range between 1st and 3rd quantile, top whisker = 1.5 * first 1st, bottom whisker = 1.5 * 3rd quantile (each of the four data quantiles contains 25% of the points). Statistical differences are indicated by different letters. **Top** depicts the sepal width (ANOVA, F = 49.16, Degrees of Freedom between species = 2, Degrees of Freedom within species = 147, P-value = 4.49 * 10-17). **Bottom** depicts petal length (Kruskal-Wallis, Chi squared = 130.41, P-value = 4.80 * 10-29).

For the remaining 17 variables, which did not meet ANOVA assumptions, non-parametric Kruskal-Wallis tests were conducted. Significant differences were identified for 16 variables (p < 0.05). Pairwise comparisons were performed using Dunn’s test with Bonferroni corrections, revealing significant groupings among species. The results, including Kruskal p-values, Chi-squared statistics, and significant pairwise comparisons, were saved in summary_kruskalwallis.csv. Annotated boxplots were generated to visualize these differences (e.g., Fig. 3).

## DISCUSSION

*Orangutan* provides a comprehensive, user-friendly framework for conducting phenotypic analyses in a comparative framework, integrating multiple statistical approaches into a single pipeline. By automating data cleaning, outlier identification and removal, allometric correction, univariate and multivariate analyses, and generation of publication-quality visualizations, *Orangutan* significantly reduces the effort and potential for error commonly encountered in species description workflows (Chan and Grismer, 2021).

Our application of *Orangutan* to empirical datasets demonstrates its utility in revealing both clear and subtle phenotypic patterns. The summary statistics function rapidly generated a detailed overview of trait variation across species (Table 1), facilitating initial assessments of central tendency and dispersion. In particular, the identification of non-overlapping diagnostic traits streamlines the process of pinpointing characters that may serve as reliable species identifiers, addressing a critical need in taxonomic practice (Balakrishnan, 2005; Cadena and Zapata, 2021). These findings underscore the value of automated range-based comparisons for expediting the discovery of diagnosis, which are often laborious to detect through visual inspection.

Optional allometric correction effectively removed size-related variation from mensural traits, enabling a focus on shape differences. Subsequent multivariate analyses—PCA and DAPC—revealed coherent species clusters in reduced-dimensional space (Fig. 2), with polygons facilitating the visual delineation of groups of interest (e.g., *A. allisoni* vs. *A. carolinensis*). The DAPC cross-classification table (Table 2) demonstrated high assignment accuracy, indicating that phenotypic data alone can robustly predict species identity when proper size correction and discriminant modeling are applied. These results support the combined use of allometric adjustment and classification-based multivariate methods for species delimitation studies.

The univariate testing framework in *Orangutan* dynamically selects parametric or non-parametric tests based on assumption checks, ensuring appropriate statistical inference for each trait. Boxplots annotated with post-hoc group letters provide clear, publication-ready visual summaries of significant differences. This flexibility enhances reproducibility by standardizing decision criteria for test selection and presentation, alleviating the burden of manual assumption testing and result formatting.

Compared to existing software, *Orangutan* uniquely combines diagnostic trait detection, size-correction workflows, and both parametric and non-parametric hypothesis testing within a cohesive R environment. Other tools may offer individual components—such as PCA visualization or basic ANOVA functions—but lack an end-to-end solution tailored specifically to the needs of systematists working on species descriptions using phenotypic data. By bridging this gap, *Orangutan* promotes more transparent, efficient, and reproducible phenotypic analyses. By automating key analytical steps and producing high-quality visual outputs, *Orangutan* enhances the democratization, reproducibility and efficiency of taxonomic research, ultimately contributing to more accessible and robust species delimitation and descriptions.

This package can provide a powerful framework for identifying distinct patterns of variation and informing comparative studies in evolutionary biology, taxonomy, and ecology.

## Data Accessibility Statement

The data to reproduce this work and software are freely and publicly available at https://github.com/metalofis/Orangutan-R.

## Author Contributions section

Javier Torres: Conceptualization, Data Curation, Formal Analysis, Investigation, Methodology, Project Administration, Resources, Software, Supervision, Validation, Visualization, Writing – Original Draft Preparation, Writing – Review & Editing

## ACKNOWLEDGEMENTS

Thanks to Mark Herr (University of Kansas) for suggestions on some analyses. Thanks to Fernando Machado Stredel (University of New Mexico), Leonardo Gonçalves and Colin Meiklejohn (University of Nebraska-Lincoln) for reviewing earlier versions of the manuscript.

## REFERENCES

Balakrishnan, R., 2005. Species Concepts, Species Boundaries and Species Identification: A View from the Tropics. Systematic Biology 54, 689–693. 10.1080/10635150590950308

Braby, M.F., Hsu, Y.-F., Lamas, G., 2024. How to describe a new species in zoology and avoid mistakes. Zoological Journal of the Linnean Society 202, zlae043.

Bro, R., Smilde, A.K., 2003. Centering and scaling in component analysis. Journal of Chemometrics 17, 16–33. 10.1002/cem.773

Cadena, C.D., Zapata, F., 2021. The genomic revolution and species delimitation in birds (and other organisms): Why phenotypes should not be overlooked. The Auk 138, ukaa069.

Chan, K.O., Grismer, L.L., 2021. A standardized and statistically defensible framework for quantitative morphological analyses in taxonomic studies. Zootaxa 5023, 293–300.

Dinno, A., 2024. dunn.test: Dunn’s Test of Multiple Comparisons Using Rank Sums.

Jombart, T., 2008. adegenet: a R package for the multivariate analysis of genetic markers. Bioinformatics 24, 1403–1405. 10.1093/bioinformatics/btn129

Kassambara, A., 2023. ggpubr: “ggplot2” Based Publication Ready Plots.

Nakagawa, S., Kar, F., O’Dea, R.E., Pick, J.L., Lagisz, M., 2017. Divide and conquer? Size adjustment with allometry and intermediate outcomes. BMC Biol 15, 107. 10.1186/s12915-017-0448-5

Neuwirth, E., 2002. RColorBrewer: ColorBrewer Palettes. 10.32614/CRAN.package.RColorBrewer

Torres, J., 2024. Gonadal Dysfunction in Abnormal-Looking Anoles Is Consistent with Hybridization and Reproductive Isolation. Ichthyology & Herpetology 112, 168–179. 10.1643/h2023065

Torres, J., Reilly, D., Nuñez-Penichet, C., Reynolds, R.G., Glor, R.E., 2025. A revision of the Anolis carolinensis subgroup supports three species in Cuba, including a new cryptic species (Squamata: Anolidae). VZ 75, 107–126. 10.3897/vz.75.e152054

Wickham, H., 2016. ggplot2: Elegant Graphics for Data Analysis.

Wickham, H., François, R., Henry, L., Müller, K., Vaughan, D., Software, P., PBC, 2023. dplyr: A Grammar of Data Manipulation.

Wickham, H., Girlich, M., 2022. tidyr: Tidy Messy Data.

